# Can the marginal male hypothesis explain spatial density variation in pinnipeds?

**DOI:** 10.1101/2025.03.24.644795

**Authors:** Koen J. van Benthem, Rebecca Nagel, Lara Maleen Beckmann, Joseph I. Hoffman, Meike J. Wittmann

## Abstract

The marginal male hypothesis is a mechanism proposed to explain gregariousness, for example in pinnipeds. Here, we explore whether this mechanism, combined with density-dependent pup survival, can also account for heterogeneity in density across colonies, as observed for example in Antarctic fur seals (*Arctocephalus gazella*). We build a discrete-time matrix model inspired by the Antarctic fur seal to investigate how differences in density between two colonies can emerge through mate choice and density-dependent pup mortality. Our model assumes a heritable male colony preference that is coupled to competitive ability, i.e. a local siring advantage. Female colony preference is modelled as an independent trait that is allowed to evolve separately. Pup survival decreases with local density. Based on numerical analysis of the model, we find slight differences in density between the colonies at equilibrium. When siring advantage is high, and the survival penalty for males with siring advantage is low, the colony that is preferred by the high-quality males actually becomes the less dense colony. Our model serves as a proof of concept that such density differences can emerge. We expect that by including additional environmental factors, future models will explain a larger part of the observed variation in density.

## Introduction

The strength and direction of sexual selection can depend on population density^1–3^. For example, in *Enchenopa* treehoppers, female preference for frequently signalling males depends on density^4^. Similarly, in sticklebacks, females that encounter few mates early in life become less choosy later in life^5^. Conversely, sexual selection may also affect patterns of local population density, for example, when sexual selection induces aggregations where males display to potential mating partners^6^. The occurrence and size of such aggregations varies considerably, even among closely related species such as pinnipeds^7^. Here, the phocid species (true seals) aggregate to a much lesser extent than the highly polygynous otariids (eared seals), with certain species like South American fur seals forming actual leks^8,9^. A number of hypotheses have been put forward to explain this variation in mating systems and the degree of aggregation, including predation avoidance, male harassment avoidance, and mate choice^7,10,11^. Here, we explore whether the effects of local density through such aggregation behaviour combined with density-dependent mortality can lead to spatial variation in density within a species.

Our study is motivated by observations of two breeding colonies of Antarctic fur seals (*Arctocephalus gazella*, Peters 1875) on Bird Island, South Georgia, that exist in close proximity. These colonies are less than 200m apart, yet have markedly different densities^12–14^. For example, between 2010 and 2016, the density of breeding females was around four times higher at the high-density colony^13^. The maintenance of this variation in density is surprising because females from both locations share the same foraging grounds and no differences have been found in quality traits such as body size and condition^12^. Simultaneously, at high density, pups are more likely to starve, be trampled or die from infection^15,16^. This negative density feedback could be expected to equalize densities between the two colonies. On the other hand, an increase in density may improve pup survival when population density is low^12,17^ (i.e. an Allee effect). The observed differences in density could also potentially be explained by the mating system. Dense female aggregations have been attributed to several factors^10^, one of which is the ‘marginal male hypothesis’^11^. Briefly, it argues that many males compete for available territories in high density breeding aggregations. If male success in this competition has a genetic component, females that leave the aggregation and spread out more evenly would risk mating with lower quality males (so-called ‘marginal males’). As a result, both sexes may favor aggregations to maximize reproductive success.

As the marginal male hypothesis revolves around female mate choice, linking mate choice to colony choice requires males to arrive in the colony first, as is the case in the Antarctic fur seal. Here, the males come ashore around a month before the females and compete to monopolise prime territories^18^. The quality of a territory is determined by the expected number of females that will arrive in that territory later to give birth and then mate^19^. A female preference may evolve if the quality of a male, including his ability to obtain a territory, is a heritable trait; however, it is unknown whether this trait is indeed heritable. However, we will assume that this necessary condition for the marginal male hypothesis is met. Females should thus prefer to mate with high-quality males in order to produce sons with higher competitive ability and reproductive success. Consequently, choosy females who favour males with high genetic quality gain fitness benefits through their offspring (which we will refer to as ‘indirect fitness benefits’). Simultaneously, an increase in female preference for breeding in a high-density colony is expected to increase the fitness of the males in that colony. Hence, a positive evolutionary feedback should emerge between the ability of a male to secure a territory in high density areas and female preference for these males. Without any costs, this could lead to runaway selection^20,21^ where both females and males develop an ever-increasing tendency to aggregate in dense patches. However, density-dependent pup mortality might limit this runaway process as aggregations become ever denser. Hence, a combination of the marginal male effect and density-dependent pup mortality could potentially explain the maintenance of spatial variation in density.

Similar indirect fitness benefits have been studied, also using a two-locus model, with one locus encoding female preference and a second locus encoding a male display trait that comes at a survival cost (e.g.^22^). In such a system, a set of mutually neutral equilibria in terms of the fraction of males that display the costly trait and the fraction of females that prefer a displaying male emerges^22,23^. As a consequence, we would expect to find stochastic drift among the different neutral equilibria. Queller (1987)^24^ elaborated on the Kirkpatrick (1982)^22^ model to show how female choosiness can lead to the maintenance of two different discrete lek sizes that are determined by male genotype. However, this model has two important limitations. First, fixed aggregation sizes for each of the two types (displaying and non-displaying males) over time are assumed, i.e. the aggregations are much smaller than the total population size and individuals can always join aggregations of their preferred size. However, in natural systems, a feedback between the fitness of certain types and the density in different areas may be expected. Second, the fitness cost of breeding at higher density is applied to the males themselves, although field data suggest that at least part of the cost is often imposed on the offspring^12^. Kirkpatrick (1982)^22^ note that such direct fitness costs to the offspring should prevent the maintenance of a polymorphism. It is, however, unclear whether a polymorphism can be maintained when the imposed survival cost on the offspring is density-dependent instead of fixed. More generally, the hypothesis that females can compensate for direct fitness losses if the surviving offspring themselves have higher fitness has also been termed the Kirkpatrick-Lande-Fisher model of runaway selection^25^, otherwise known as the ‘sexy son hypothesis’. The occurrence of such runaway selection is not limited to two-locus systems, but has also been demonstrated using a quantitative genetics model, where female preference and the male display trait are genetically correlated^26^.

Here we evaluate whether a combination of indirect fitness benefits through competitive sons and density-dependent pup mortality could explain the observed spatial heterogeneity in density in pinnipeds, using the Antarctic fur seal as a motivating example. So far, this is only a verbal model and in a system like this with the potential for complex direct and indirect effects, a quantitative model is needed to determine whether or not the proposed mechanisms could lead to the observed variation in density^27^. We therefore use a matrix model that tracks genotype distributions over time in a two colony setup for a diploid, two-sex population, and perturb this model to disable either the direct fitness effects of density or indirect fitness effects in order to understand the role of these effects as well as their interactions. We expect that, if indirect fitness effects are disabled by allowing females to mate with males from either colony, population densities should even out and male quality should go to a single maximum value.

## Methods

Our model describes a population of Antarctic fur seals whose choice for one of two colonies depends upon both sex and genotype. We assume two autosomal diploid loci with two alleles each, leading to a total of nine possible (multilocus) genotypes that are tracked through time (Fig. 1). Each locus is expressed in one sex only and the loci are inherited independently.

**Figure 1.**
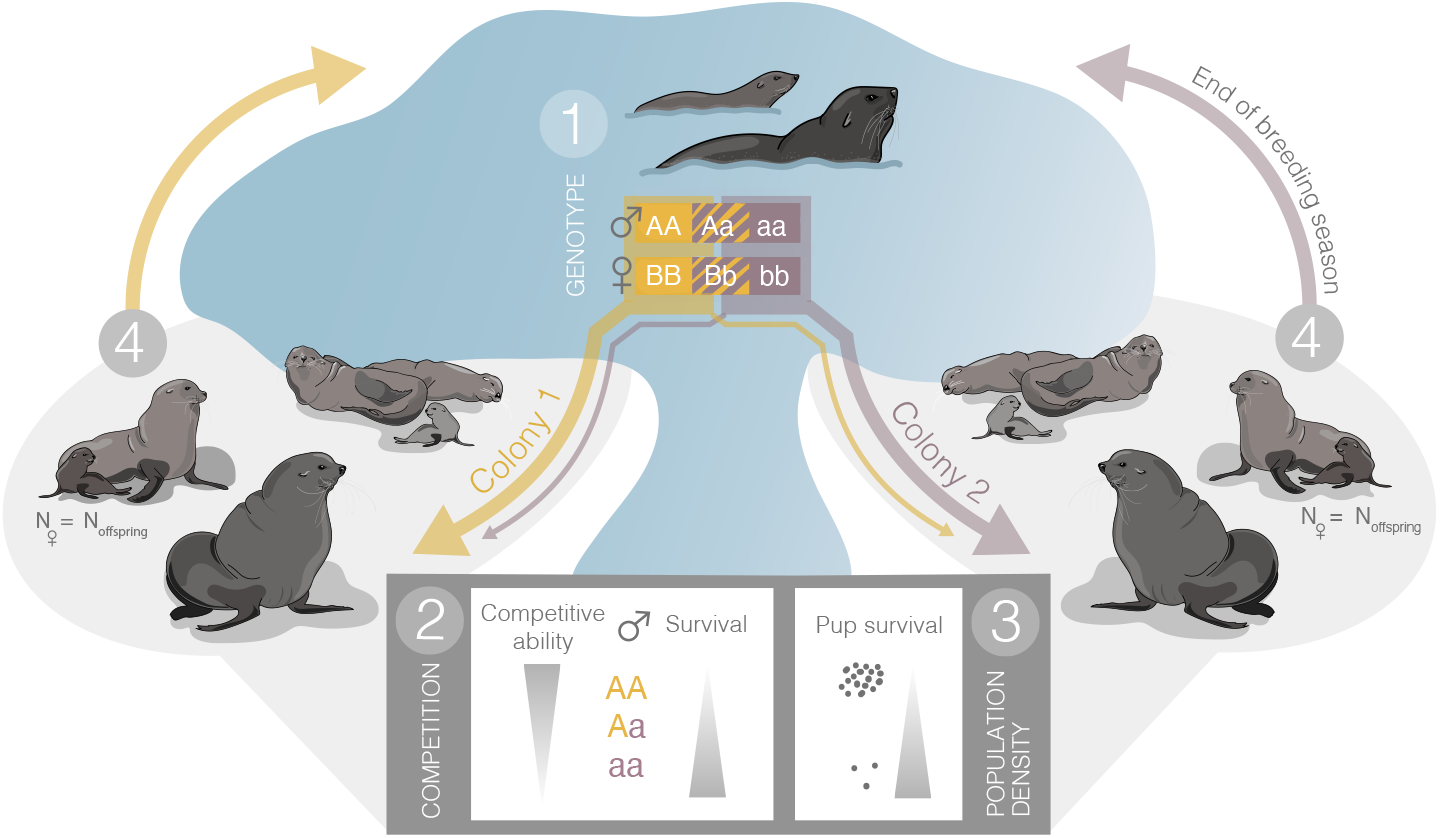
Schematic depiction of our model. 1. Fur seal genotype is coded by two loci. The first locus with A and a alleles is expressed in males, and determines their colony preference, with AA males preferring to go to colony 1. The second locus with alleles B and b analogously determines the site preference of females. 2. Besides colony preference, the genotypes further affect competitive ability and survival in the males. 3. Colony choice may affect local population densities, which in turn affects pup survival. 4. At the end of the time step (i.e. season) the overall population density is regulated.

The first locus, with alleles A and a, codes for the competitive ability of males; the AA and Aa males have an increased chance of siring offspring, with a factor *P*_1_ and 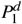, respectively, compared to aa males in the same colony. Here 0 ≤ *d* ≤ 1 refers to the dominance: when *d* = 0.5, in any given colony the reproductive output of AA males (*f*_*AA*_) relative to that of Aa males is the same as reproductive output of Aa relative to that of aa males. This local increase in mating success comes at a cost, however, and these males have a reduced survival of 1 − *δ* for AA males and (1 − *δ*)^*d*^ for Aa males. Male site preference is also determined by the first locus, with an AA male going to colony 1 1 −Γ_*m*_ and an aa male to colony 2 with probability 1 −Γ_*m*_. Here Γ_*m*_ represents the chance of a homozygous male going to the non-preferred patch and it should have a value ≤ 0.5. Heterozygous males have an intermediate site preference depending on the dominance 0 ≤ *d*_2,*m*_ ≤ 1 and go to colony 1 with probability (1 −Γ_*m*_)*d*_2,*m*_ + Γ_*m*_(1 − *d*_2,*m*_) = Γ_*m*_ + *d*_2,*m*_(1 − 2Γ_*m*_).

The second locus, with alleles B and b, determines the site preference of a female. Analogous to the first locus, females with BB go to colony 1 with probability 1 −Γ _*f*_ and those with bb go to colony 2 with probability 1 −Γ _*f*_ . Heterozygous females have an intermediate site preference and go to colony 1 with probability Γ _*f*_ + *d*_2, *f*_ (1 − 2Γ _*f*_).

The total population can be described by a population vector 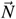:

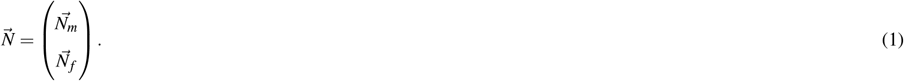

Here, the sex-specific abundance vectors for males and females, respectively, are defined as:

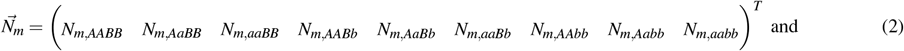

and

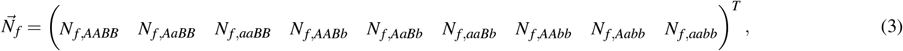

where *T* denotes the transpose. In total, the nine different genotypes, combined with 2 different sexes, yield a total of 18 discrete classes to keep track of. The number of individuals in each of the 18 classes in colony *i*, 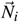 are not tracked explicitly, but instead they can be calculated at any time from the overall genotype distribution and the corresponding site preferences:

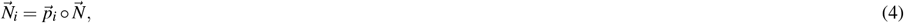

with ° the element-wise product and *-p*_*i*_ the probabilities for each of the classes to go to colony *i*, as described above. In particular:

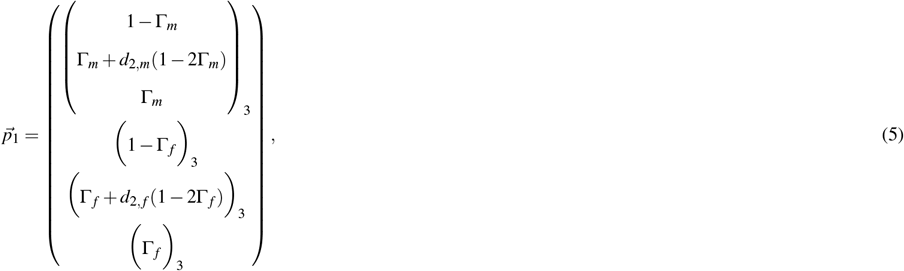

where the subscripts refer to repeated elements, such that

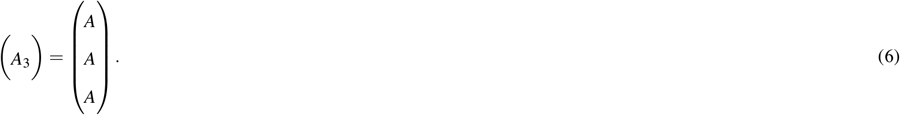

Because we model only two colonies:

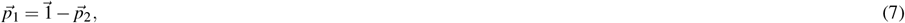

and hence:

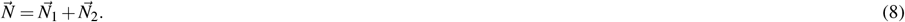

The offspring distribution is determined from the male and female distributions. At each time step, the number of offspring born at a patch is equal to the number of females in that patch. This choice is a simplification as in nature reproductive rates for females are known to be around 0.75 on average^19^. Because the population is renormalized after each time step, we do not expect the rate to qualitatively affect the outcomes. For the offspring, an equal sex ratio is assumed, and the offspring genotype distribution depends on the distributions of maternal and paternal genotypes in the colony. Compared to males with an aa genotype, males with an AA genotype have a *P*_1_ times higher chance of siring an offspring and males with an Aa genotype 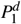 times higher chance. In the model, this is achieved by multiplying the relative abundance of AA males by *P*_1_ and Aa males by 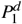and then renormalizing the male distribution. From these two genotype distributions, the offspring genotype distribution is calculated, assuming Mendelian inheritance and genetically unlinked loci. Potential covariation between alleles across the two different loci is accounted for by explicitly evaluating all combinations of male and female genotype classes and their expected offspring distributions.

The survival rate of offspring depends on the density of the colony where they are born. The higher the density, the greater is the chance that a pup dies^15,16^. As we do not know the exact extent to which male versus female density affect pup survival, we use a weighted average of male and female density, with a weighting factor *β* . The survival rate of offspring in colony *i* is given by:

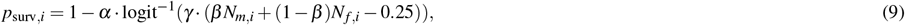

with *α* the relative strength of density regulation and *N*_*f,i*_ the number of females in colony *i*:

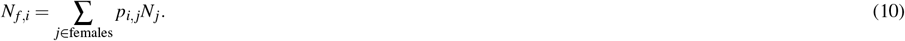

and *N*_*m,i*_ analogously corresponding to the number of males in colony 1. The strength of density regulation (*α*) can vary between 0 and 1 and determines the maximum survival penalty due to trampling. Fig. 2 illustrates the shape of Eq. 9. Note that the 0.25 in Eq. 9 corresponds to an even number of males and females in both colonies: the population is normalized at each step (i.e. total density of 1) and thus if there are equal numbers of males and females, and these numbers are identical in both colonies, the density of each sex in each colony would be 0.25.

**Figure 2.**
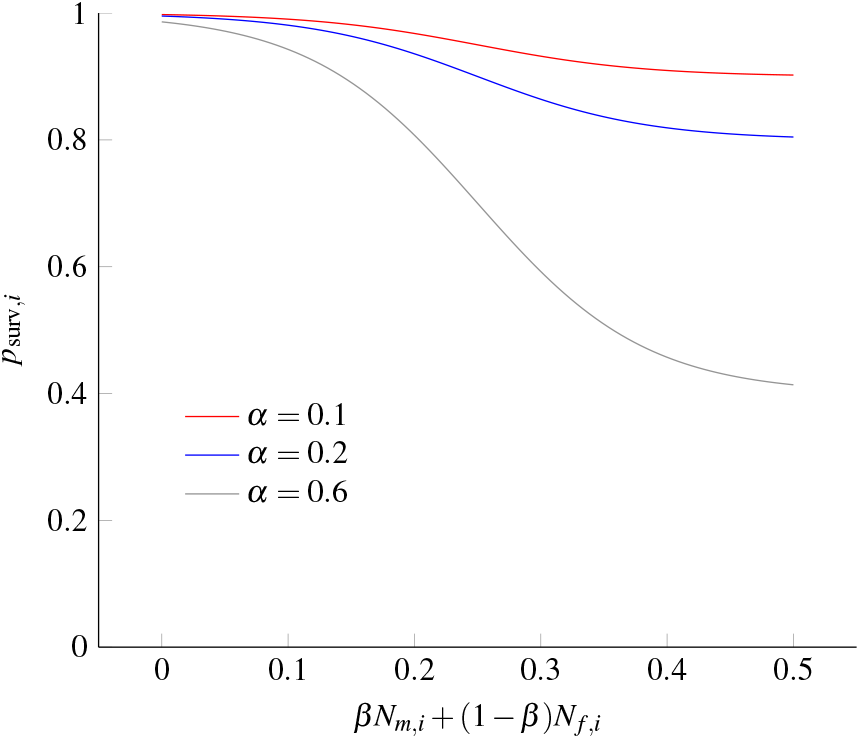
Illustration of pup mortality as a function of the weighted average density in the colony, *β N*_*m,i*_ + (1 −*β*)*N*_*f,i*_ (see Eq. 9), for different values of *α*.

Finally, global density regulation is imposed by normalizing the distribution at every time step.

Maternal colony choice affects female long-term fitness through two pathways: density-driven pup mortality and the local genetic composition of potential mating partners. To assess the impact of the second pathway on population dynamics, we also ran the model under random mating. In this scenario, fathers were selected independently of their colony when determining the genetic distribution of offspring, while density-dependent pup mortality remained locally regulated. By comparing these runs to those where colony choice influenced mate availability, we were able to disentangle the effects of the two pathways.

## Results

Before systematically changing the parameters, we first present two example simulations in Fig. 3. In the first model run (Fig. 3A), males and females eventually became unevenly distributed across the two colonies. This is an indirect consequence of the lower survival of AA males, because AA males that preferably settle in colony 1 have reduced survival, leading to lower male abundance in colony 1. Starting from an even distribution of females, this leads to a lower overall density in colony 1 and, as a consequence, increased pup survival in that colony. After these initial dynamics, females develop a slight preference to move to colony 1. As a consequence, the relative fitness of AA males increases, by increasing their per-capita fecundity, and the overall density across the two colonies evens out, although colony 1 remains more female-biased. Whether the total abundance exactly evens out depends on the precise parameter settings. Fig. 3B illustrates the case where males suffer a low survival penalty and a high siring advantage compared to 3A, leading to colony 2 becoming the more populated colony overall, due to there being more females in that colony.

**Figure 3.**
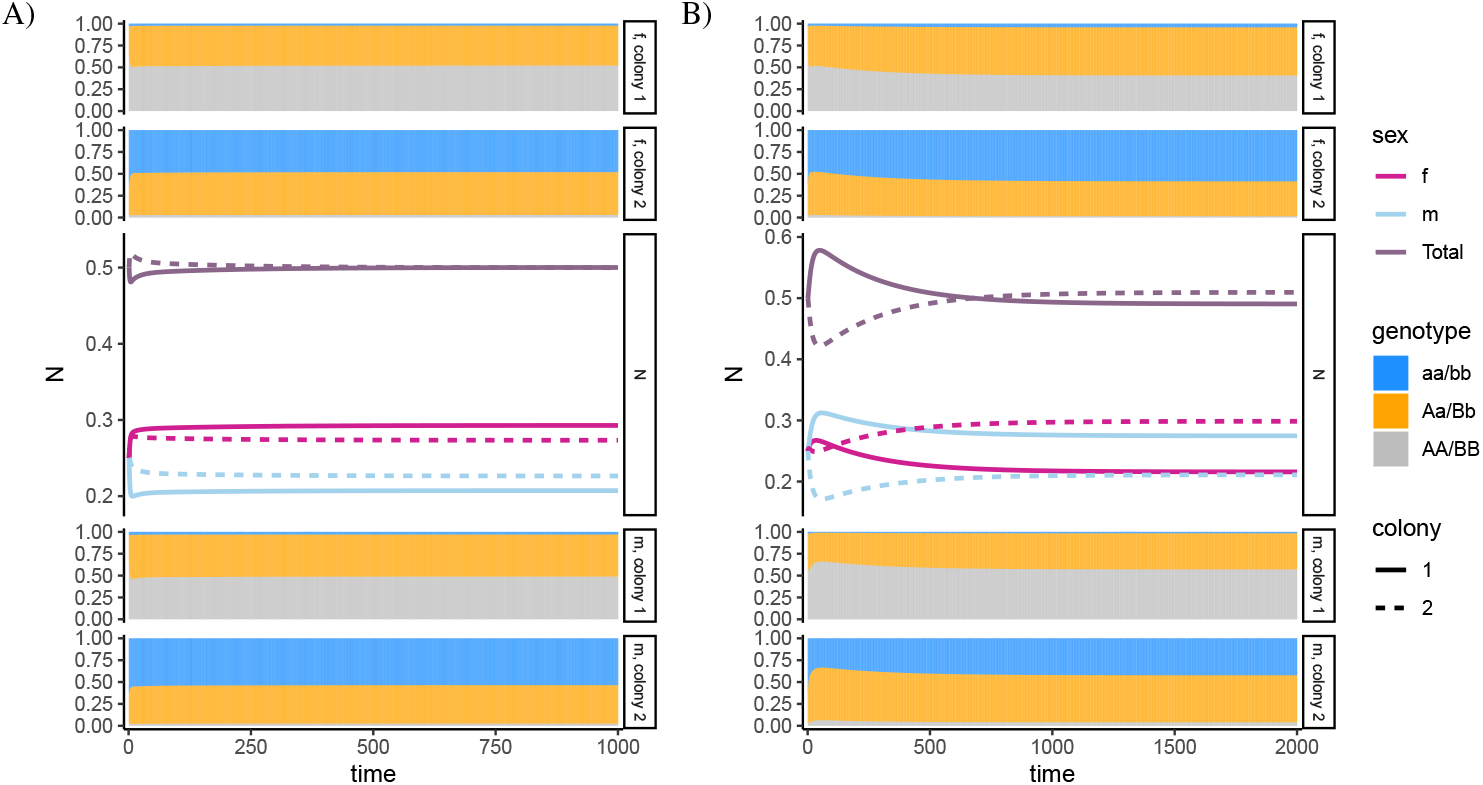
Two example time series with different parameter settings are depicted in panels A and B respectively. In both cases, initially, the population vector was uniform. From top to bottom the five panels contain: 1) Female genotype (bb, bB and BB) frequencies over time for colony one, 2) female genotype frequencies in colony two, 3) Total male and female abundance over time (middle panels), 4) Male genotype (aa, aA and AA) frequencies over time for colony one, and 5) Male genotype frequencies for colony two. Parameter values: *α* = 0.05, *γ* = 15, *β* = 0.5, *d* = 0.5, *d*_2,*m*_ = *d*_2, *f*_ = 0.5, Γ_*m*_ = Γ _*f*_ = 0.05, and A) *P*_1_ = 1.8, *δ* = 0.3, and B) *P*_1_ = 3, *δ* = 0.05.

We now turn to systematically varying siring advantage, survival penalty and density-dependent pup mortality, in order to investigate how these affect the densities of the two colonies. We first focus on the siring advantage of AA and Aa males and their survival penalty (Fig. 4 upper row). As a general consequence of the direct fitness effects of density, males and females tend to develop opposing colony preferences. For example, when AA males suffer a strong survival penalty and have a weak monopoly on paternity (top left corner of the panels in Fig. 4A top row), females prefer colony 1, while males tend to go more often to colony 2. Our model therefore predicts that females prefer to go to colony 1, even if the males therein have no competitive advantage (i.e. *P*_1_ = 1), but do suffer a strong survival penalty, and are thus of lower quality, in contrast to our initial expectations. Consequently, the number of females per male in colony 1 is highest when the AA males have a relatively small siring advantage combined with low survival (Fig. 4B, top left panel, top left corner). This leads to higher per-capita fecundity in colony 1 for the males and thereby facilitates slightly higher female densities, but the overall density is still highest in colony 2. Colony 1 has a higher overall density when the siring advantage of AA males is high and their survival is low (upper right corner). Overall it seems that the fitness advantage due to increased local competitive strength is typically counterbalanced by an emerging lower per-capita male fecundity due to a lower female to male ratio (in colony 1 female to male ratio decreases with siring advantage of AA males, while it increases in colony 2, see Fig. 4B).

**Figure 4.**
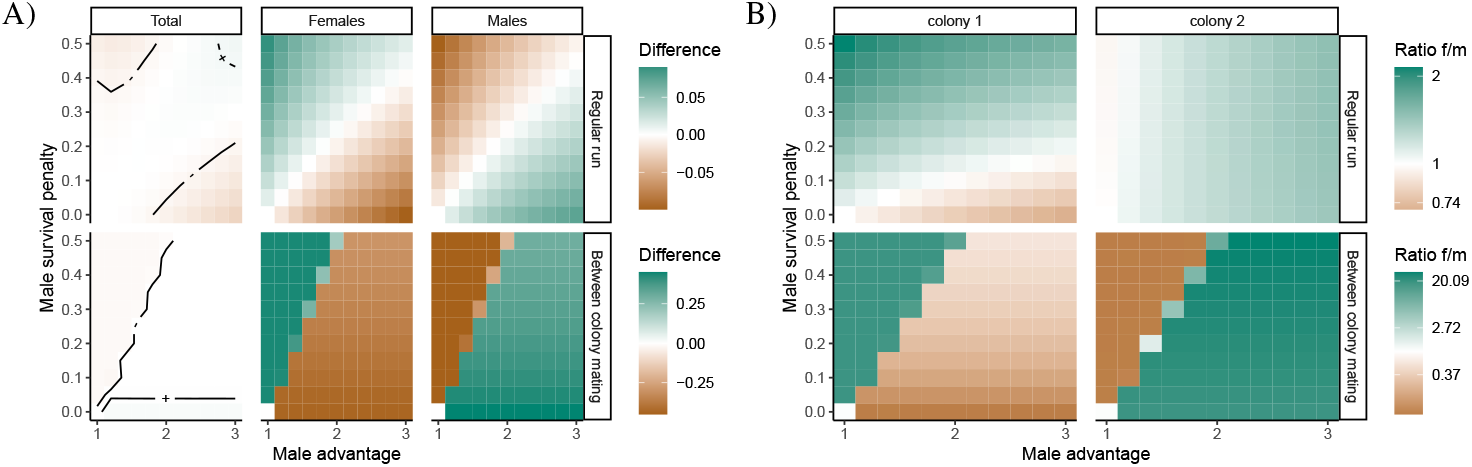
The effect of male advantage (*P*_1_) and male survival penalty (*δ*) after 10^4^ time steps. A) Difference in abundance between colony 1 and 2, with positive numbers indicating more individuals in colony 1. Contour lines in the left panel correspond to a difference of 0.005 (+) or -0.005 (-). B) The resulting female/male ratio in each colony. The result for each unique combination of *P*_1_ and *δ* was obtained by averaging over 15 replicate simulation runs, each of which was initialized with a different randomly drawn initial population distribution. The top row in both panels correspond to the model run as described in the methods, while the bottom row corresponds to a perturbed run in which females can mate with males in the other colony too. All other parameters are as described in the legend of Fig. 3.

In a perturbed version of the model where the indirect fitness benefits are disabled by allowing females to choose a mating partner from either colony, the sex-specific patterns are similar, but overall density effects are different (Fig. 4 bottom row).

Here, when the AA males are relatively fit (i.e., when their mating advantage is high and their survival penalty is low, Fig. 4, lower row, below the diagonal), the AA males dominate, which can be seen from the differences in male density between the colonies. These differences are relatively large compared to the null expectation of 0.25 males and 0.25 female density in each colony, when both alleles are equally common and individuals distribute themselves uniformly. The females in this model do not rely on colony choice for their mating partner and therefore evolve to go to colony 2, avoiding the high male density of colony 1. This leads to an equalization of the overall density across the colonies. In contrast, when the AA males are of relatively low quality compared to the aa males, the opposite pattern emerges, with colony 2 becoming male dominated, and colony 1 being female dominated (4, bottom row, above the diagonal). Importantly, when disabling the effect of female patch preference on male per capita fitness, the overall higher abundance that we observe for colony 1 in the top right corners of the panels of Fig. 4 is much weaker. This supports the above explanation that the higher density of colony 1 in this region of the parameter space depends on effects mediated by local mate availability, which in turn affect the per capita fecundity of the males.

Fig. 5 shows how the population dynamics depend on the density effect on offspring. The left-most columns within the top panels in Fig. 5, where the density-dependent pup mortality disappears completely, exhibit stochastic patterns that depend on the initial conditions of the system. However, as soon as density-dependent mortality is non-zero, the system reaches equilibrium abundances that are independent of the initial conditions. Furthermore, these abundances hardly vary with increasing density-dependent mortality. This general pattern remains unaltered when females are able to choose their mate from either colony (Fig. 5, bottom row). However, this is not the case when there is no survival penalty for the offspring. In this case, the females tend to distribute evenly on average across trials, while the males display site preference for colony 1 when the survival penalty is low and for colony 2 when it is high. Furthermore, as with our initial results, sex-specific differences in

**Figure 5.**
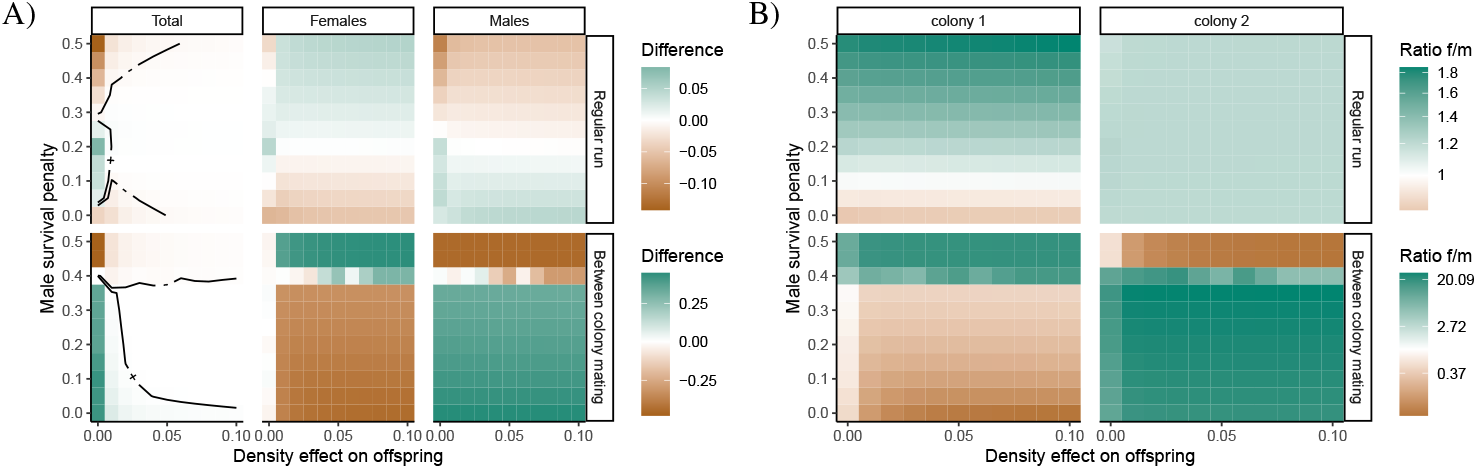
The effect of offspring survival penalty (*α*) and male survival penalty (*δ*) on A) the difference in abundance between the two colonies after 10^4^ time steps, with positive numbers indicating more individuals in colony 1, with contour lines in the left-most panel corresponding to a difference of 0.005 (+) or -0.005 (-), and B) the female/male ratio in each colony The result for each combination of *α* and *δ* was obtained by averaging over 15 replicate simulation runs, each of which was initialized with a different randomly drawn initial population vector. All other parameters as in Fig. 3.

density become far more pronounced when mate choice is not restricted to one’s own colony, and eventually this leads to each colony containing almost exclusively only one sex (Fig. 5B, bottom row). The similarity in patterns between the case where mate choice is colony-restricted and the case where it is not suggests that population dynamics in this region of the parameter space are largely driven by direct density dependence rather than by indirect second-generation fitness effects through genetic quality of the offspring.

Finally, we explored how our results are affected by the genetic link between patch choice and quality (Fig. 6). We show that as the uncertainty in male colony choice (Γ_*m*_) increases, while keeping the siring advantage constant, females prefer to go to colony 2 - to counter high male densities in colony 1, as even in colony 2 there is a good chance of mating with an AA male. Hence, when male colony choice becomes completely random, the female preference again disappears. Colony 1 only becomes the denser colony when female and male colony choice are both relatively uncertain. This is driven directly by the evening out of the females, due to their absence of colony preference, while AA males benefit from being spread evenly across colonies, giving them access to mating partners in both colonies. When male colony choice becomes completely random, the total abundance evens out. It is also important to note that when females can freely choose their mating partner from either colony, the pattern changes much more strongly compared to the other scenarios. Now, females prefer to go to colony 2 when colony uncertainty is relatively low in both sexes, to counter the high male density in colony 1. However, this effect disappears as colony choice becomes less certain, as the direct effect of density disappears and there is no difference between the colonies in terms of the choice of available partners.

**Figure 6.**
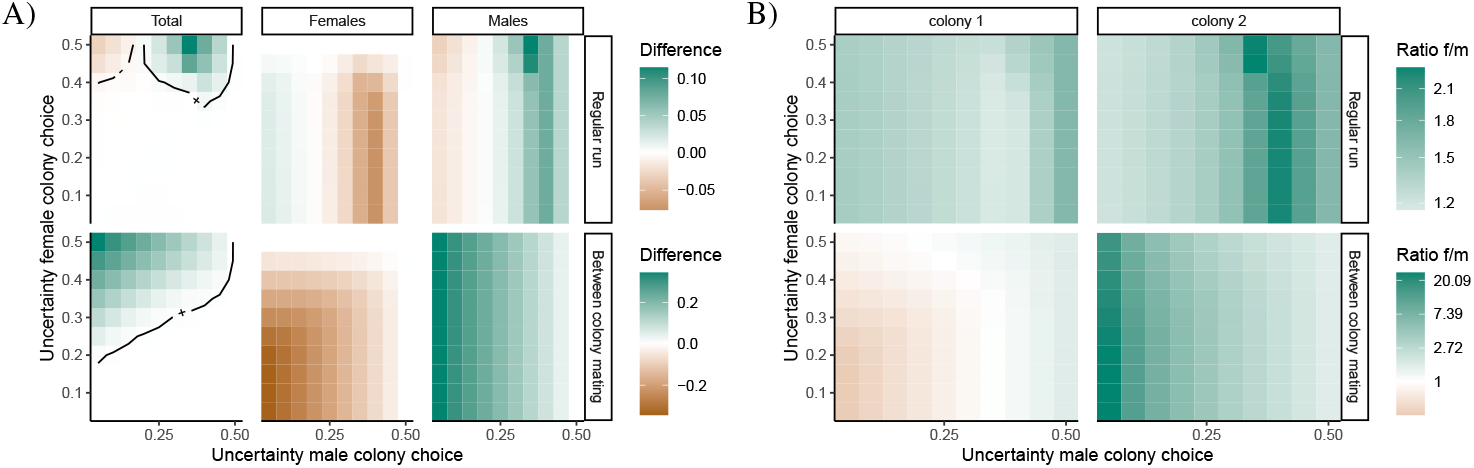
The effect of uncertainty in male (Γ_*m*_) and female colony choice (Γ _*f*_) on A) the difference in abundance between colony 1 and 2, with positive numbers indicating more individuals on island 1, and contour lines in the left most panel corresponding to a difference of 0.005 (+) or -0.005 (-), and B) the resulting female/male ratio on each island. Shown are the situation after 10^4^ time steps, averaged over 15 replicates, each with a randomly drawn initial population vector. All other parameters as in Fig. 3.

## Discussion

Our proof-of-concept model shows that sexual selection coupled with local density-dependent fitness can produce differences in population density across space and maintain variation in male quality and female patch preference. Mechanistically, this arises through a combination of direct fitness effects of colony choice and indirect fitness benefits for females, whose colony choice affects the genetic quality of their offspring. Our model was inspired by two naturally occurring populations of Antarctic fur seals, but our results can potentially extend to other systems in which density-dependent offspring mortality and aggregation behaviour exist.

Our results show that emerging differences in density between the colonies are highest when colony choice is relatively uncertain (weak colony preference, see Fig. 6. When colony preferences are strong (uncertainty in colony choice of 0.05), the differences in density are generally small (i.e. not exceeding 5% of the overall population size, see Fig. 4A). A higher overall density in colony 1 occurs when the AA males suffer from a relatively high survival penalty and have a strong local siring advantage (top right corner in the panels in Fig. 4 top row). Here, males seem to be slightly overrepresented in colony 2, while colony 1 is mainly female-dominated. When the siring advantage is low and the survival penalty is high, or when the survival penalty is low and the siring advantage is high, density is highest in colony 2. An example time series for this latter scenario is shown in 3B, where colony 2 becomes the more populated colony, with a large number of females in that colony. This appears to be caused by AA males being of such high quality, both in terms of survival and siring advantage, that colony 1 initially becomes overly crowded (Fig. 3B). The females in turn start avoiding the relatively densely populated colony 1, leading to reduced per-capita fecundity in the AA males by inducing a male-biased sex ratio in colony 1 and further reducing the selective advantage for females to go to colony 1. The finding that females evolve to avoid colony 1 and thereby AA males when these males are of higher quality (i.e. higher survival and high siring advantage) is contrary to our initial expectation.

### Coupling of genetic quality and patch preference

In the model, genetic variation in both male and female traits is maintained. This is achieved through the genetic coupling of local siring success to patch preference. The extent to which the mating advantage of AA males can play out depends on the strength of local competition in their colony as well as on the local availability of females: if most females prefer to go to the second colony, for example to avoid density-dependent pup mortality, the per-capita fitness of AA males might actually be lower than that of aa males. Density-dependent pup mortality limits the total number of individuals in either colony, while the sex ratio determines the per-capita fecundity of the males. When mating occurs between individuals from different colonies (e.g. Fig. 4A bottom row) females can mate with high quality males, without having to raise their pups in the respective colony. This leads to strong segregation, with males and females inhabiting different colonies, thereby reducing the effect of density-dependent pup mortality. For example, when siring advantage is high and survival penalty low for AA males, we end up with strongly male biased population in colony 1 (AA males) and a female biased population in colony 2 (bb females, Fig. 4B, bottom right corner of the panels). As a consequence, genetic variation in male quality and female patch preference is much lower. When mating can occur between individuals from different colonies (Fig. 4B bottom row), genetic variation in males disappears completely and only AA males remain (results not shown).

We do not know the mechanism through which colony preference is inherited. Two main mechanisms that could contribute to colony preference are that the preference for a specific (type of) colony is heritable, or that animals genetically inherit an intrinsic tendency to return to their specific birth place (natal site philopatry). Although both mechanisms could be relevant in the actual population, we here only chose the former mechanism for the link between genotype and colony choice, because this mechanism allows us to track preference for specific colonies more directly. Developing a model focusing on natal site philopatry and comparing the two would be a valuable future research direction. If a heritable preference for a specific colony exists, offspring are likely to have a similar colony preference to their parents and one would expect to observe natal philopatry. In female Antarctic fur seals, natal philopatry has been observed on a fine scale for locations within a colony^28^. Males that are resighted across multiple years also tend to hold territories in the same locations within a colony^29^. If philopatry in both males and females is strong across colonies, it reduces gene flow between them. We observe this in our model, where the genetically determined colony preference leads to a linkage disequilibrium between the A and B loci (see Fig. S1.1). Hence, females preferring to go to either colony also tend to carry the gene for their sons to go to the same colony. As a consequence, genetically differentiated populations might emerge if enough time elapses, colony choice is highly reliable, and local populations are small or exposed to strong local selection pressures. However, microsatellite data from the two focal colonies of Antarctic fur seals used as our motivating example suggest that they are not genetically differentiated^30^, although population genetic structure can be found on a larger geographic scale^31,32^. This suggests either that colony preference is less heritable than we assumed, or that some of the other conditions for differentiation have not been met.

### Spatial heterogeneity

Our model functions as a proof-of-concept that slight spatial heterogeneity in population density can emerge without fundamental differences between the two colonies. The underlying mechanism is the coupling of male local siring success to colony preference. Hence, we invoke processes at different spatial scales: local competition among males for mating partners and local density-dependent pup mortality, combined with the colony preference of the females in the overall population and global density regulation. Indeed, it is known that multiscale processes can lead to the emergence of heterogeneity in density, in particular in the presence of an Allee effect (see e.g.,^33,34^). However, these models differ from the model presented here, in the sense that they do not consider both sexes simultaneously and hence cannot take into account the effects of female density on male fitness. In continuous space, clusters of individuals can also emerge over ecological timescales as a consequence of biased density-dependent movement^35^. Although our model is defined in discrete space, it bears similarity to biased movement because patch preference is included as an evolvable trait.

The formation of clusters can also be explained following the classical lek system. The larger an aggregation of males, the more females it should attract because these females would have a higher chance of finding a high-quality male when there are more males present^6^. This mechanism requires female mate choice to occur within a colony, which has been documented in the Antarctic fur seal^36^. An assessment of male quality could, for example, be based on the density of males in the colony, since high density signals the capacity of a male to hold a territory even in a place where the potential for conflict is very high. However, investigating female mate choice and the assessment of partner quality and in particular the heritability of these traits will probably require a quantitative genetic analysis of male quality, the heritability thereof, and how it relates to environmental factors such as density, through the usage of an animal model (see e.g.,^37^).

Although our model does find heterogeneity in density, the magnitude of the difference in density is relatively small. The empirically observed differences in density are thus likely determined by additional factors such as underlying environmental factors^38^. For example, in Antarctic fur seals, females tend to gather in closer proximity on shingle beaches compared to boulder beaches^39^. However, topographical differences alone are unlikely to explain the observed difference in density, as similar differences in density have been observed between colonies that are share a more similar topography^15^ (i.e. both being surrounded by cliffs). Our model shows one potential mechanism through which heterogeneity in density may emerge in an otherwise homogeneous landscape.

## Conclusions

We have shown that indirect fitness effects combined with density-dependent pup mortality can lead to variation in density. Counterintuitively, our results show that if high-quality males have both high survival and high siring advantage, their preferred colony will become the less dense colony. When survival is low, and siring advantage is high, we find the more intuitive result that the beach that is preferred by the high-quality males becomes the denser beach. In the future, our model could be extended to explore the interplay of the intrinsic factors studied here and extrinsic environmental factors, such as predation, in determining variation in local population density.

## Author contributions

MW, JH and KvB conceived the ideas and designed the methodology. KvB led the analyses and the writing of the manuscript. LB provided the schematic depiction of the model. All authors contributed to interpretation of the results. All authors reviewed and edited the manuscript.

## Acknowledgements

This research was funded by the German Research Foundation (DFG) as part of the SFB TRR 212 (NC3)-project numbers 316099922, 396774617, and 396782288.

