## Supplementary material for "Can the marginal male hypothesis explain spatial density variation in pinnipeds?": SI S1

### S1 Linkage disequilibrium

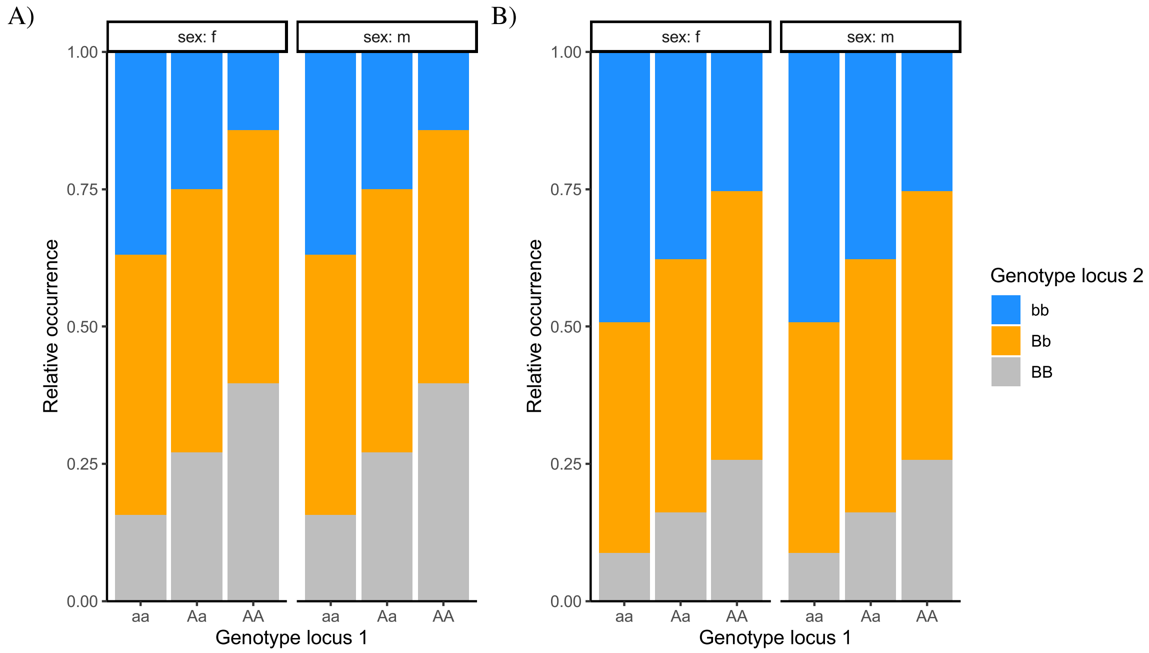


**Figure S1.1.** Coupling between the two loci at the last time step of the two example time series from Fig. 3. Shown is the relative occurrence of the b-genotype within each a-genotype.
